# Insights on the GPCR helix 8 solution structure and orientation using a neurotensin receptor 1 peptide

**DOI:** 10.1101/2024.01.31.578299

**Authors:** James B. Bower, Scott A. Robson, Joshua J. Ziarek

## Abstract

G-protein coupled receptors (GPCRs) are the largest class of membrane proteins in the human genome with high pharmaceutical relevance and implications to human health. These receptors share a prevalent architecture of seven transmembrane helices followed by an intracellular, amphipathic helix 8 (H8) and a disordered C-terminus. Technological advancements have led to over 1000 receptor structures in the last two decades, yet frequently H8 and the C-tail are conformationally heterogeneous or altogether absent. Here we synthesize a peptide comprising the neurotensin receptor 1 (NTS1) H8 and C-terminus (H8-Ctail) to investigate its structural stability, conformational dynamics and orientation in the presence of detergent and phospholipid micelles, which mimic the membrane. Circular dichroism (CD) and nuclear magnetic resonance (NMR) measurements confirm that zwitterionic 1,2-diheptanoyl-sn-glycero-3-phosphocholine is a potent stabilizer of H8 structure, whereas the commonly-used branched detergent lauryl maltose neopentyl glycol (LMNG) is unable to completely stabilize the helix – even at amounts four orders of magnitude greater than its critical micellar concentration. We then used NMR spectroscopy to assign the backbone chemical shifts. A series of temperature and lipid titrations were used to define the H8 boundaries as F376-R392 from chemical shift perturbations, changes in resonance intensity, and chemical-shift derived phi/psi angles. Finally, the H8 azimuthal and tilt angles, defining the helix orientation relative of the membrane normal were measured using paramagnetic relaxation enhancement (PRE) NMR. Taken together, our studies reveal the H8C-tail region is sensitive to membrane physicochemical properties and is capable of more adaptive behavior than previously suggested by static structural techniques.

## STATEMENT

G protein-coupled receptors are arguably one of the most pharmacologically relevant receptors in the human body. Understanding the molecular nature of these receptors will serve to expand upon their therapeutic impact. The present study helps to outline the structural and dynamic properties of a understudied region under solution conditions. This work simultaneously expands on the possible NMR methodologies that can be applied to these challenging systems.

## INTRODUCTION

G protein-coupled receptors (GPCRs) comprise the largest protein family in eukaryotes – accounting for 3-4% of the human-genome.[1] With over a third of FDA-approved drugs targeting GPCRs, it follows that these receptors possess considerable significance in human physiology and pathology.[2] They share a similar architecture of seven transmembrane helices with a majority also possessing an amphipathic helix 8 (H8) followed by a C-terminal tail (C-tail). Amongst Class A GPCRs, H8 possesses a semiconserved F(R/K)xx(F/L)xxx(L/F) motif while the C-tail shows very little to no sequence identity between receptor families.[4] Previous studies highlights the importance of H8 in receptor topogenesis, trafficking and surface expression [5-8]. For example, H8 and Ctail mutation/truncation in the bradykinin B_2_ receptor (B_2_R) attenuates receptor trafficking to the plasma membrane while also altering interactions with G proteins, β-arrestins and GRKs. Similarly, a handful of works highlight the importance of Helix 8 and the Ctail in G-protein[13-15] and arrestin interactions [11, 16-18].

Over the past few decades, advances in X-ray crystallography and cryo-electron microscopy (cryo-EM) structural biology have led to the generation of 172 unique receptor structures with 133 of these being Class A GPCRs at the time of writing. [3] Despite the significance of H8 and the C-tail in receptor topology and function, it has not been a major focus within these structural studies apart from a few investigations on rhodopsin and µ-opioid receptors[9-12]. Although this is not for a lack of effort, this regions is frequently truncated to achieve structure determination or altogether absent in electron density [4]. The neurotensin receptor 1 (NTS1) serves as a prime example with only 11 of the 23 rat NTS1 crystal and cryo-EM structures exhibiting a structured H8 helix [19-21]; the helical boundaries vary between models and, in all cases, presents the highest B-factor across the receptor (Figure 1a and S1).

**Figure 1.**
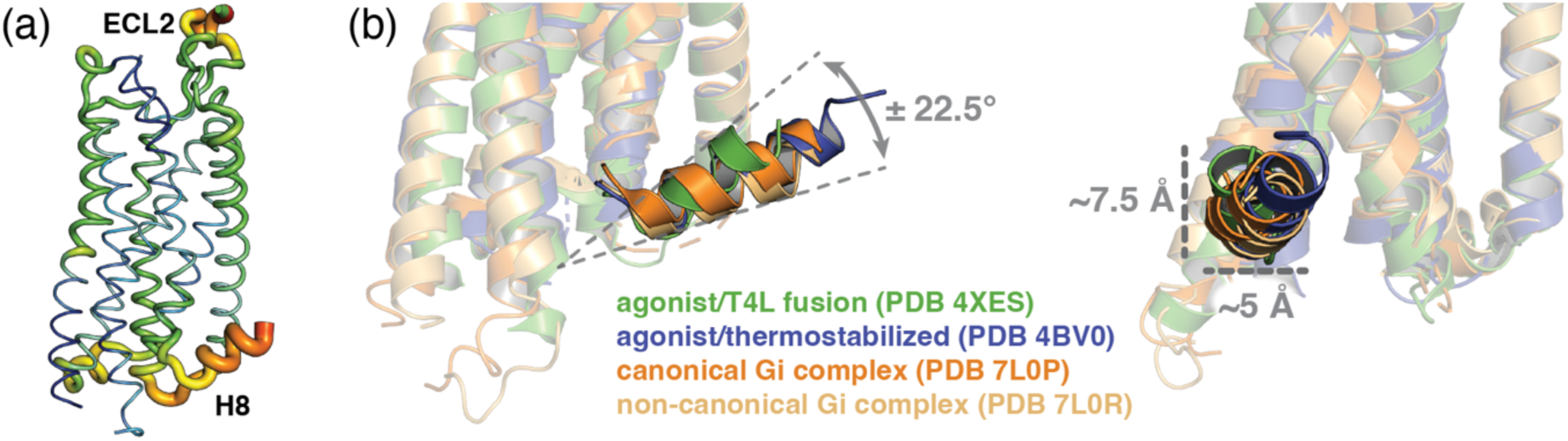
NTS1 structures feature variable helix 8 lengths and orientations. a.) PyMOL B-factor putty display of a representative agonist-bound rat NTS1 (PDB 4XES) crystal structure. b.) Overlay of rat NTS1 structures: agonist-bound/T4L fusion (PDB 4XES; green), agonist-bound/thermostabilized (PDB 4BV0; blue), canonical Gi complex (PDB 7L0P; orange), and non-canonical Gi complex (PDB 7L0R; light orange). Approximate angle between most extreme helix 8 positions of 4XES and 7L0R is shown. Approximate scale around helix 8 positions is shown.

The NTS1 H8 tilt angle varies >20° that produce rigid-body translations up to 7.5 Å as measured at the terminal residues (Figures 1b and S2). The receptor construct and sample preparation likely play a role in this variability, but may also reflect the physicochemical differences between detergent micelles, lipid nanodiscs, and lipidic cubic phase membrane mimetics [22]. It’s also worth noting that all NTS1 structures include thermostabilizing mutations, C-terminal/intracellular loop 3 (ICL3) truncations and/or the addition of stabilizing molecules such as T4-lysozyme (T4L) [20, 23, 24], DARPin [25], or an antibody fragment[26].

Here, we used a synthetic rat NTS1 peptide comprising the putative H8 region through the C-terminus to characterize the structural stability of this region in the solution state. First, we used circular dichroism (CD) to test the impact that temperature and membrane mimetic composition have on H8 secondary structure. We show that this construct requires a detergent or lipid micelle to stabilize the helical structure, although the degree of helicity depends on the nature of the micelle. Next, we produced an isotopically labeled peptide to measure residue specific secondary structure by NMR. Analysis of chemical shift perturbations, resonance intensity, and chemical shift-derived backbone torsion angles indicate the most stably-structured H8 boundaries converge to residues F376-R392. Dynamic NMR experiments reveal intermediate timescale motions in residues adjacent to these boundaries, including one of the putative palmitoylation sites[27], which we hypothesize reflects transient helix (un)folding. Finally, paramagnetic relaxation enhancement (PRE) NMR was used to determine the H8 tilt and axial rotation angle relative to the lipid micelle.

## RESULTS

### Phospholipids and detergents stabilize NTS1 H8 to variable extents

We synthesized a peptide, herein termed pNTS1(H8-Ctail), corresponding to the putative rNTS1 helix 8 through the C-terminus, residues S373^7.56^-Y424 (superscript refers to Ballesteros-Weinstein nomenclature)[28]. The C386 and C388 palmitoylation sites[27] were substituted with serine to prevent disulfide formation. We collected far-UV CD spectra of pNTS1(H8-Ctail) in aqueous buffer at increasing temperatures from 5 °C to 90 °C (Figure 2a). These spectra revealed a generally unstructured peptide characterized by a strong negative signal around 200 nm at lower temperatures. Interestingly, with increasing temperature the 200 nm signal became weaker (less negative) and the 222 nm intensity strengthened (more negative). These spectral changes are consistent with induction of PPII helical character at increasing temperature [29]. An isodichroic point at ∼207 nm indicates this is a two-state conformational equilibrium.

**Figure 2.**
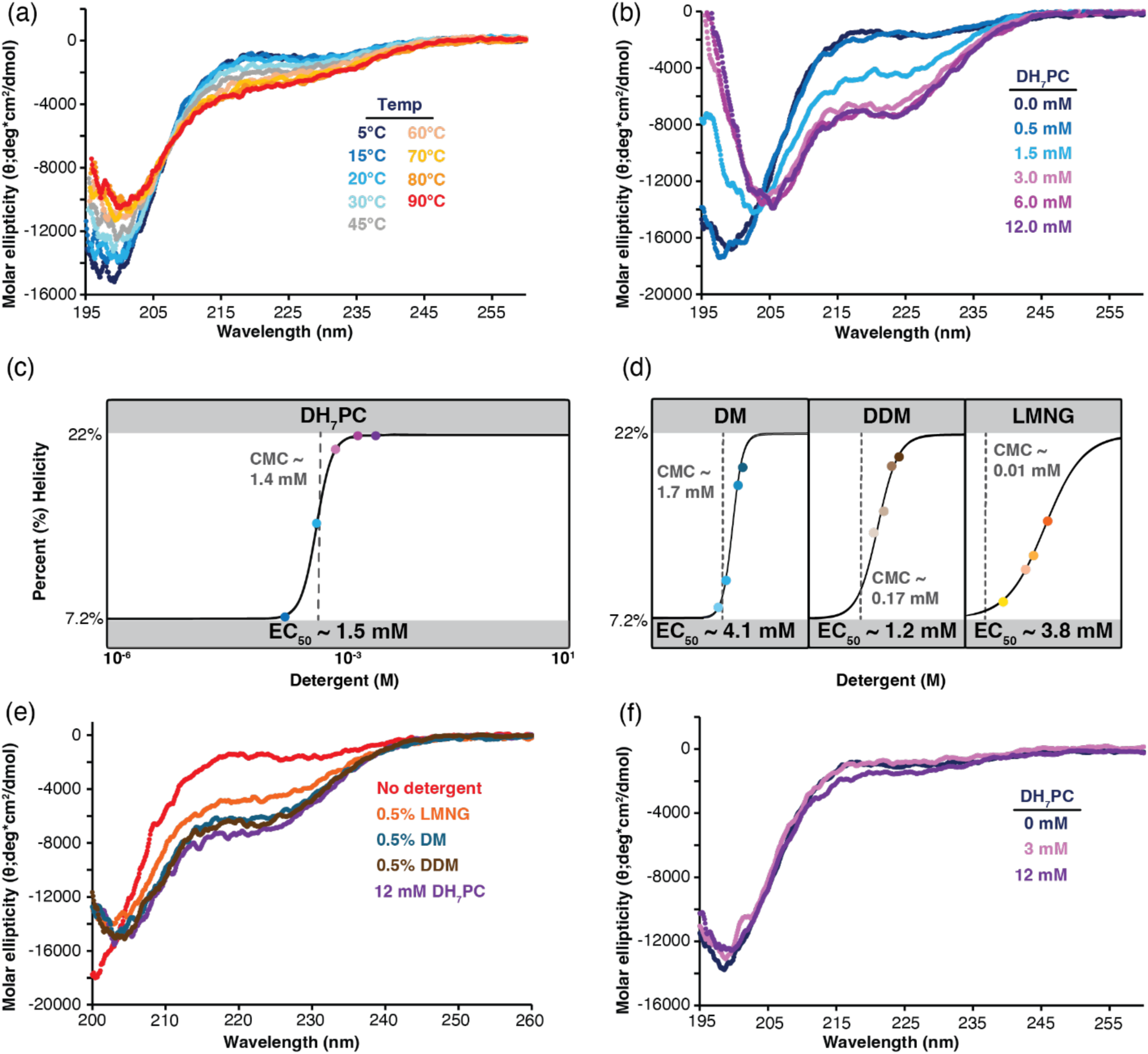
Membrane mimetics induce variable amounts of helical character in the NTS1 peptides. a.) Overlay of pNTS1(H8-Ctail) temperature series CD spectra. An isodichroic point is observed at ∼207 nm indicative of a two-state equilibrium. b.) Overlay of pNTS1(H8-Ctail) CD spectra collected at 20 °C with increasing DH_7_PC concentrations. An isodichroic point is observed at ∼205 nm indicative of a two-state equilibrium. c.) The percent helicity of pNTS1(H8-Ctail) induced by DH_7_PC at 20 °C was calculated from the 222 nm molar ellipticity using the Lifson-Roig-based helix-coil model [31, 32]. The percent helicity was then fit to a four-parameter model to estimate the EC_50_; the CMC is indicated by a gray dashed line. d.) The percent helicity was calculated and fitted for NTS1(H8-Ctail) with increasing concentration of DM, DDM, or LMNG detergent at 20 °C. Each CMC is indicated by a gray dashed line. e.) CD spectral overlay of pNTS1(H8-Ctail) in highest detergent concentrations of DM, DDM, LMNG, and DH_7_PC at 20 °C. f.) CD spectra overlay of pNTS1(C-tail) in increasing DH_7_PC concentrations at 20 °C. All spectra were collected using 35 μM pNTS1(H8-Ctail) in aqueous buffer with 10 mM NaPi (pH 6.8).

Next, we titrated in diheptanoyl-sn-glycero-3-phosphocholine (DH_7_PC) to test if phospholipids affect the secondary structure of pNTS1(H8-Ctail). Whereas lipid concentrations below the critical micellar concentration (CMC ∼ 1.4 mM) had little to no effect on the pNTS1(H8-Ctail) CD spectrum, helix formation is clearly initiated at 1.5 mM DH_7_PC as indicated by an increasingly negative 222 nm signal (Figure 2b). Greater lipid concentrations stabilized the helical architecture until it saturated between 3-6 mM; this suggests a 1:1 peptide:micelle stoichiometry assuming DH_7_PC has an aggregation number of 50.[30] Again, an isodichroic point at ∼205 nm indicates a two-state folding equilibrium. The molar ellipticity was fitted to the Lifson-Roig-based helix-coil model from Baldwin et. al that assumes ellipticity at 222 nm is linearly related to mean helix content (Figure 2c) [31, 32]. Given the 35 μM pNTS1(H8-Ctail) concentration, these results indicate that pNTS1(H8-Ctail) is 7.3% helical in the absence of lipid. DH_7_PC induces 22.2% helical character, or approximately 11.5 residues, which is consistent with bioinformatics predictions that H8 comprises 15 amino acids.[33] Next, we tested if three other commonly used non-ionic detergents (n-decyl-beta-maltoside, DM; n-dodecyl-beta-maltoside, DDM; and lauryl maltose neopentyl glycol, LMNG) are sufficient to similarly re-organize the pNTS1(H8-Ctail) architecture. Little to no helicity was observed below their respective CMCs, but both DM and DDM produced substantial helix formation at five-fold CMC (Figures 2d,e and S3a,b). Surprisingly, LMNG induced very little helicity until concentrations exceeded ten times the CMC (Figures 2d,e and S3c). We calculated the percent helicity for each maltoside-derived detergent concentration, as above. These values were then fitted to a four-parameter dose-response model constraining the total pNTS1(H8-Ctail) helical content in the absence and presence of detergent to 7.3% and 22.2%, respectively (Figure 2d). We estimate that 31% (w/v) LMNG, or ∼31,000X its CMC, would be required to stabilize the helical structure 90% of the time.

### Phospholipids do not promote helicity in the C-tail region

Alignment of the rNTS1 sequence with other class A receptors predicts that residues 373-388 form H8 with the remaining C-terminus as a disordered tail [33]. Yet, published X-ray crystallographic and cryo-EM NTS1 structures feature a variety of helical boundaries with electron densities typically terminating near residues 383-385 [19-21]. Spin-label electron paramagnetic resonance (EPR) spectroscopy experiments indicated the helix comprised residues 374-393, although no labels were tested beyond these two extremes [4]. To test whether the helicity observed in pNTS1(H8-Ctail) originated from the putative disordered C-tail, we repeated our CD measurements on another synthetic peptide, herein termed pNTS1(Ctail), constituting residues A385^8.59^-Y424. pNTS1(Ctail) CD spectra were collected in aqueous buffer at increasing temperatures from 5 °C to 90 °C (Figure S3d). As observed above for pNTS1(H8-Ctail), the spectra indicate a generally disordered peptide with a characteristic strong negative signal around 200 nm; increasing temperature pushed the two-state equilibrium toward the PPII helix. DH_7_PC did not substantially remodel the CD spectra with a strong negative 200 nm signal persisting even at 12 mM lipid – more than eight times the CMC (Figure 2f).

### Assignment of pNTS1(H8-Ctail) NMR resonances

Next, we developed a recombinant *E. coli* expression system to isotopically-label pNTS1(H8-Ctail) for solution NMR spectroscopy. This construct comprises rNTS1 residues 373-424 with two amino acid modifications relative to the synthetic pNTS1(H8-Ctail) construct used in the CD studies above. We engineered a non-canonical 3C cleavage site to remove a C-terminal GFP-fusion, which necessitated a Y424F mutation and addition of Q425 [34]. ^15^N- and ^13^C-heteronuclear single quantum coherence (HSQC) spectra of [*U*-^15^N,^13^C]-pNTS1(H8-Ctail) were well-resolved at 5 °C without membrane-mimetics; concentration-dependent spectral overlays did not exhibit any signs of aggregation/oligomerization (data not shown). Approximately 98% of expected backbone amide and 100% of expected Cα/Cβ resonances were assigned from conventional 3D HNCA, HN(CO)CA, HNCO, HN(CA)CO, (H)CC(CO)NH, and H(CCCO)NH triple resonance experiments (Figure 3a). Secondary structure was predicted from the N, H_N_, Cα, Cβ, CO, Hα, and Hβ chemical shift values using TALOS-N [35] and, consistent with the CD results above, the peptide is disordered in the absence of lipid (data not shown). We collected a temperature series of ^15^N- and ^13^C-HSQCs to transfer assignments from 5 °C to 35 °C by visual inspection. The resonances universally traverse linear trajectories in fast-intermediate exchange as a function of temperature (data not shown) suggesting the perturbations reflect altered solvent exchange kinetics, rather than a local structural change, which is consistent with the CD temperature series.

**Figure 3.**
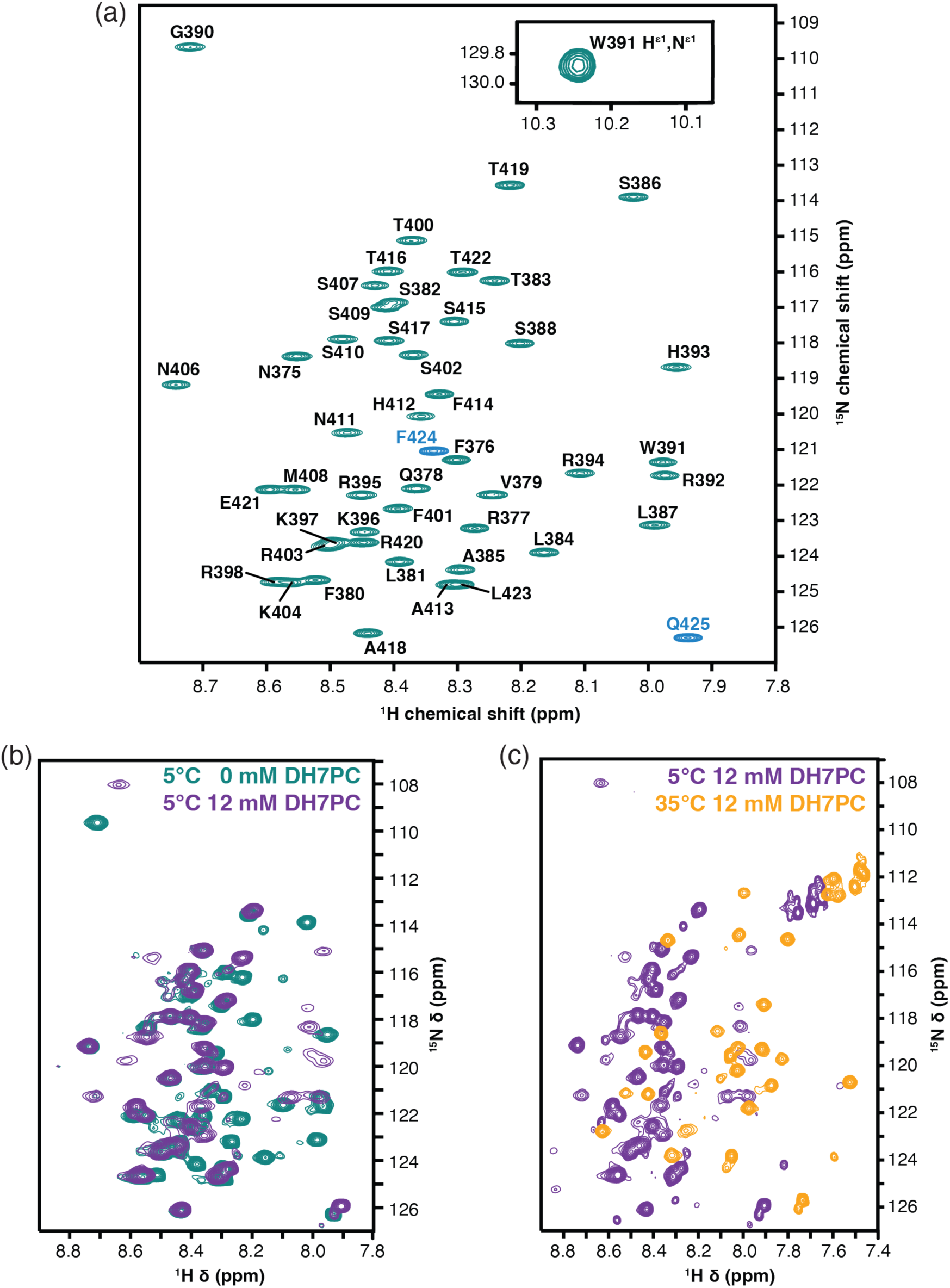
Temperature and/or lipid alter the ^15^N-HSQC spectra of pNTS1(H8-Ctail). a.) Assigned ^15^N HSQC spectra of [*U*-^15^N,^13^C]-pNTS1(H8-Ctail) at 5 °C. The last two residues of the peptide were engineered as part of the 3C protease cleavage site (blue). b.) ^15^N-HSQC spectral overlay of [*U*-^15^N,^13^C]-pNTS1(H8-Ctail) in the absence (teal) and presence (purple) of (d26)-DH_7_PC at 5 °C. c.) ^15^N-HSQC spectral overlay of [*U*-^15^N,^13^C]-pNTS1(H8-Ctail) in 12 mM (d26)-DH_7_PC at 5 °C (purple) and 35 °C (orange). All spectra were collected on 450 μM [*U*-^15^N,^13^C]-pNTS1(H8-Ctail).

### NMR reveals specific interactions of helix 8 and the C-tail with the DH_7_PC micelle

We next sought to determine specific structural boundaries of the DH_7_PC-induced helix. First, we titrated [*U*-^15^N,^13^C]-pNTS1(H8-Ctail) with 0, 0.5, 1.5, 3, 6, 10, and 12 mM tail-deuterated (d26)-DH_7_PC at 5 °C (Figures 3b and 4a). Beginning at 1.5 mM (d26)-DH_7_PC, F376-R394 ^1^H/^15^N amide resonances undergo some permutation of slow, intermediate or fast chemical exchange on the NMR timescale. If we assume the chemical shift perturbations report on a two-state equilibrium, then the slowly exchanging residues suggests a lower limit for the exchange rate (k_ex_) of 6 s^-1^ [36]. The spectral perturbations saturated at ∼12 mM (d26)-DH_7_PC, which corresponds to an approximately 1:1 peptide:micelle stoichiometry similar to CD experiments. A temperature series of ^15^N- and ^13^C-HSQC spectra were collected at 12 mM (d26)-DH_7_PC to transfer assignments from 5 °C to 35 °C by visual inspection; again, we observed substantial chemical shift perturbations across the range of timescales from slow to fast exchange (Figure 4b). Overall, N-terminal resonances gain intensity as temperature increases while C-terminal peaks simultaneously disappear (Figures 3c and 4c). The simplest explanation for these changes in the chemical exchange timescale is that N-terminal residues are bound to a faster tumbling micelle and/or the rate of micelle associate/dissociation has increased. To confirm assignments of the slowly exchanging resonances at 35 °C, we recollected HNCA, HNCO, (H)CC(CO)NH, and H(CCCO)NH spectra in 12 mM (d26)-DH_7_PC. Complete assignments were not possible for resonances undergoing intermediate timescale chemical exchange. TALOS-N predictions were successfully generated at 35 °C and 12 mM (d26)-DH_7_PC for residues 373-400, 404, 414, 415, and 418-425 (Figure 4d). Taken together, the chemical shift perturbations, intensity changes and TALOS-N predictions all support a structured H8 helix from F376-R386 followed by a weaker helix between 387-392, most likely due to the helix breaking proline at P389 (Figure 4).

**Figure 4.**
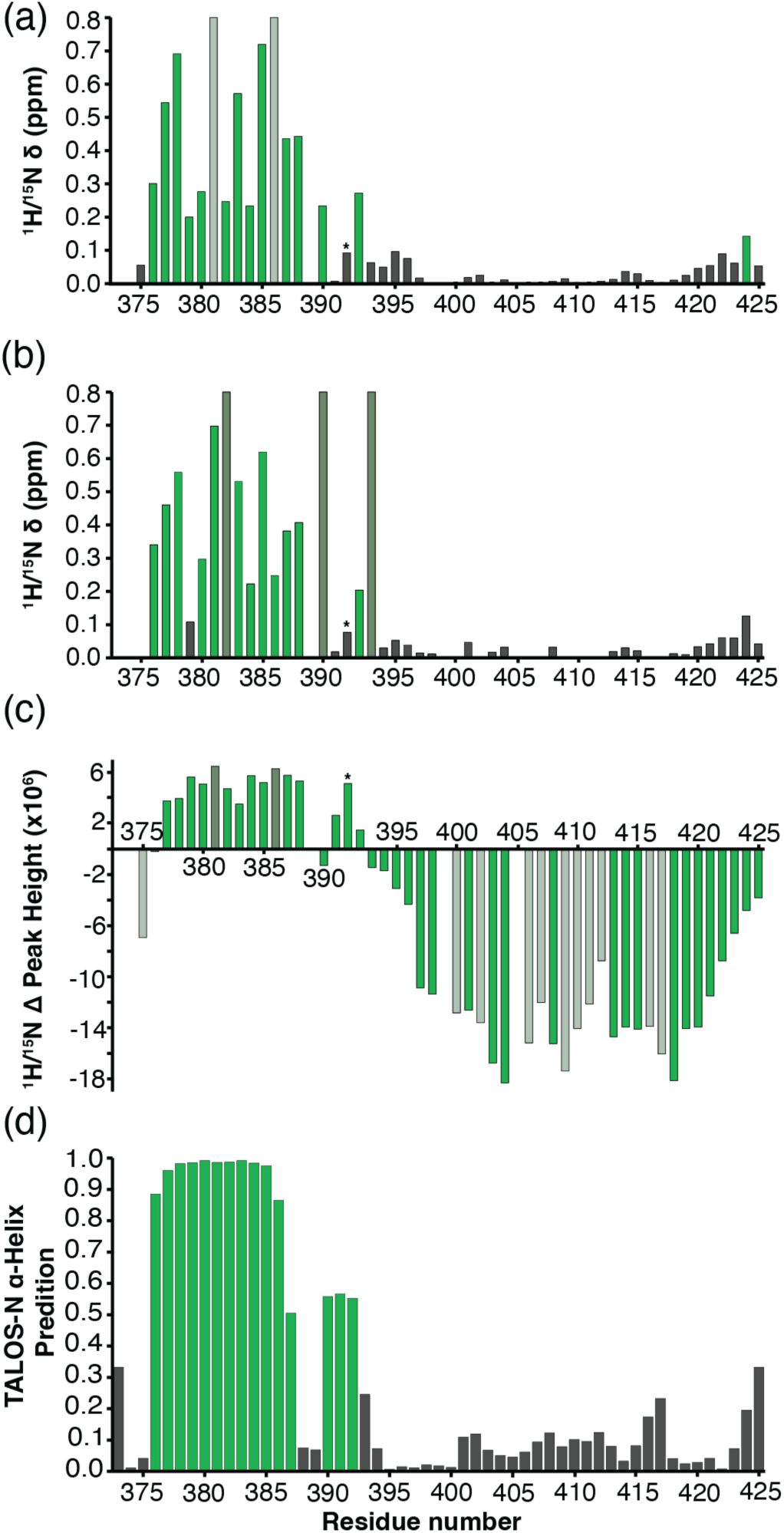
Chemical shift and peak height perturbations define the H8 helical boundaries. ^1^H/^15^N combined chemical shift perturbation of [*U*-^15^N,^13^C]-pNTS1(H8-Ctail) induced by 12 mM (d26)-DH_7_PC at (a.) 5 °C and (b.) 35 °C. c.) Change in [*U*-^15^N,^13^C]-pNTS1(H8-Ctail) ^1^H/^15^N peak height from 5 °C to 35 °C in the presence of 12 mM (d26)-DH_7_PC. In panels a-c, perturbations above threshold are green, peaks that disappear or appear are set to the largest perturbation value then colored light or dark green, respectively. d.) TALOS-N secondary structure prediction of estimated alpha helix probability for each residue with 12 mM (d26)-DH_7_PC at 35 °C. Probabilities > 0.5 are colored green. All spectra were collected on 450 μM [*U*-^15^N,^13^C]-pNTS1(H8-Ctail). The asterisks indicate W391 H^ε1^/N^ε1^.

### NMR relaxation predicts a stable helix in the presence of DH_7_PC micelle

To confirm the stability of pNTS1(H8-Ctail) helical boundaries, we measured ^15^N longitudinal (T_1_) and transverse (T_2_) relaxation times at 5 °C and 35 °C in the presence of 12 mM (d26)-DH_7_PC. Regardless of temperature, residues N375-R394 have distinct dynamic properties, supporting a discrete architecture from the remaining peptide. This region displays longer ^15^N T_1_ and shorter ^15^N T_2_ values at 5 °C with shorter ^15^N T_1_ and ^15^N T_2_ values at 35 °C, compared to the C-tail region (Figure S4a-d). Individual T_1_ and T_2_ values are difficult to interpret without an estimate of the overall rotational correlation time (τ_c_); therefore, we approximated the τ_c_ of amenable residues from their ^15^N T_1_/T_2_ ratio assuming a spherical Stokes-Einstein particle using the approach of Kay and colleagues.[37] The τ_c_ values at both temperatures indicate a slower molecular tumbling rate for the helical region compared to the C-tail (Figure 5a,b). Interestingly, at 5 °C the calculated τ_c_ values for residues N374-R392 follow an apparent inverted-U curve (Figure 5a) with the central portion of the helix having an ostensibly lengthier tumbling time – suggesting differential structural stability across this region. Such an inverted-U curve does not exist in the τ_c_ estimates for the helical region at 35 °C (Figure 5b).

**Figure 5.**
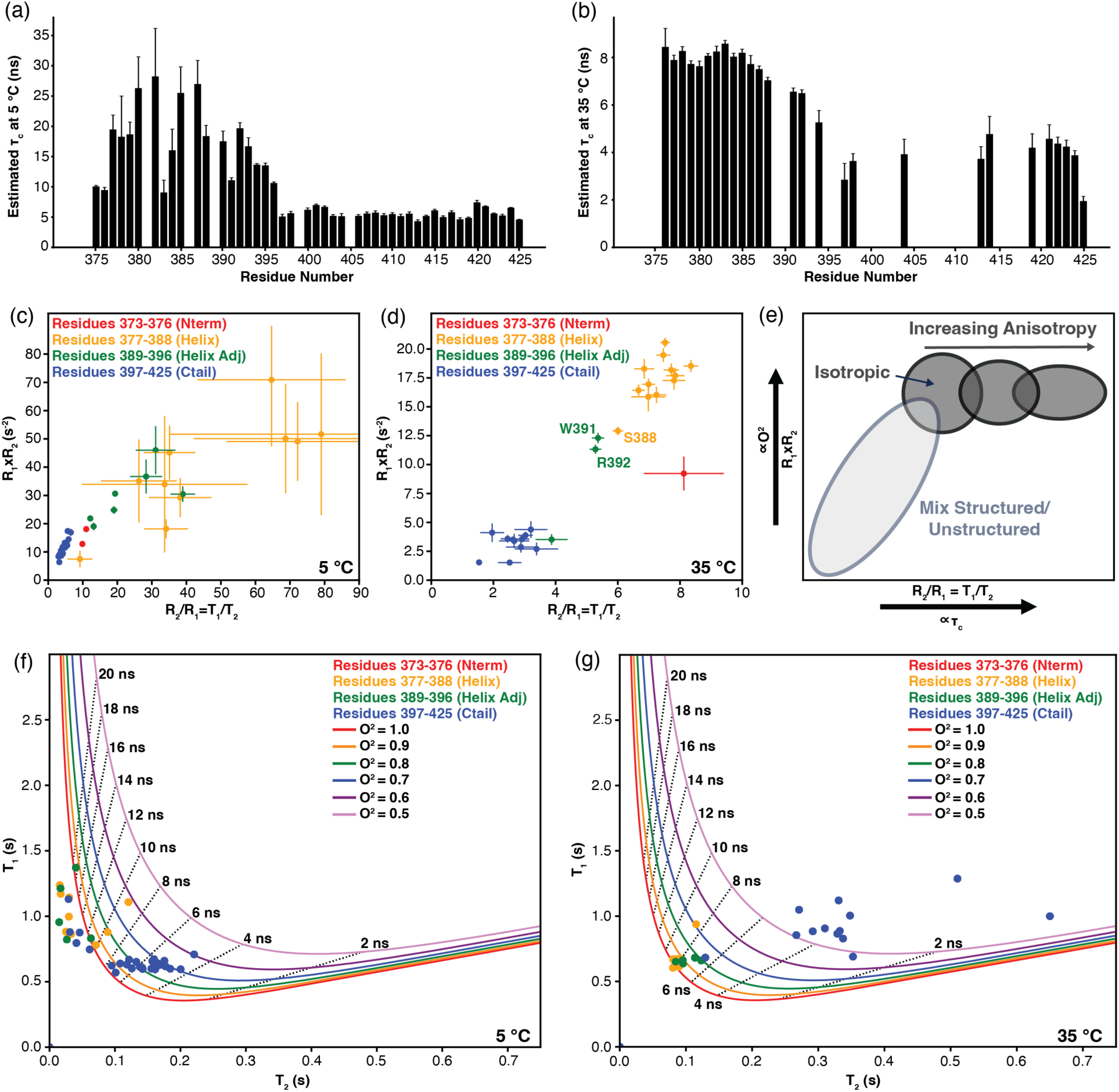
Analysis of pNTS1(H8-Ctail) T1 and T2 relaxation in presence of DH_7_PC. a.) The pNTS1(H8-Ctail) T_1_/T_2_ values were used to predict the overall rotational correlation time (τ_c_) in 12 mM (d26)-DH_7_PC at (a.) 5 °C and (b.) 35 °C. pNTS1(H8-Ctail) R_1_×R_2_ values are plotted against R_2_/R_1_ values (c.) 5 °C and (d.) at 35 °C. e.) Illustration of clustering patterns for R_1_×R_2_ vs R_2_/R_1_ plots depending on measured isotropy or anisotropy [38]. Experimentally measured T_1_ and T_2_ relaxation times for pNTS1(H8-Ctail) residues are plotted with model-free simulated T_1_ and T_2_ curves at (f.) 5 °C and (g.) 35 °C.

Processes such as binding to the slower reorienting micelle, anisotropic molecular tumbling, and/or μs-ms timescale chemical exchange generally shorten T_2_ resulting in an artificially higher T_1_/T_2_ ratio [37]. To reduce the number of factors potentially complicating the interpretation of our ^15^N relaxation data, we calculated the R_1_×R_2_ (T_1_^-1^×T_2_^-1^) product [38], which is insensitive to anisotropic molecular tumbling and serves as a proxy for the generalized order parameter (O^2^). O^2^ characterizes the degree of information on the NH bond vector orientation lost to sub-τ_c_ timescale dynamical processes, or more simply the magnitude of ps-ns timescale dynamics [38]. The R_1_×R_2_ values for residues N375-R394, at 5 and 35 °C, remain substantially elevated with an inverted-U curve distribution (Figure S4e,f). Interestingly, the inverted-U curve distribution now appears for the helix at both 5 and 35 °C, although less dramatically for 35 °C.

We then plotted R_1_×R_2_ (T_1_^-1^×T_2_^-1^; proxy for O^2^) against R_2_/R_1_ (i.e. T_1_/T_2_; proxy for τ_c_) (Figures 5c,d) [38]. The pNTS1(H8-Ctail) plots at both temperatures depict a correlated distribution with N-terminal and C-terminal residues clustered in the upper-right and lower-left quadrants, respectively (Figures 5c,d). Our interpretation of this correlation is that high order and long rotational diffusion times are a feature of structured sections, while low order and short diffusion times are associated with unstructured portions. That is, regions like the C-tail, which appear to have low R_1_×R_2_ values (low order parameters), also have faster apparent tumbling times (lower R_2_/R_1_) because their diffusion is independent of the micelle or globular protein; conversely, where the H8-Ctail interacts with the micelle, we suspect rigid structure and longer tumbling times. As shown in Figure 5e, the distribution of R_1_xR_2_ versus R_2_/R_1_ values from an isotropically-tumbling folded proteins would be expected to produce a circular-shape that stretches horizontally with increasing anisotropy [38]. Whereas the H8-Ctail peptide appears isotropic at 35 °C, which would be predicted if its motions are dominated by a nearly spherical micelle, at 5 °C the R_1_xR_2_ values plateau with increasing R_2_/R_1_ ratios suggesting an anisotropic structured state (Figure 5c). However, chemical exchange could also explain this result.

The magnitude of chemical exchange contributions can be estimated for resonances that substantially deviate from the average R_1_×R_2_ product [38]. Since the high variance across N- and C-terminal regions reduced our confidence in estimating an overall average R_1_×R_2_ value, we simulated the maximum R_1_×R_2_ at 600 MHz magnetic field strength over a range of O^2^ and τ_c_ values (Figure S5) [38]. At 600 MHz, R_1_×R_2_ should not exceed ∼20 s^-2^ (Figure S5) at the highest order limit of O^2^ = 1.0. Yet, at 5 °C residues R377-G390 are at 30-50 s^-2^ with A385 >70 s^-2^ (Figures 5c), which significantly exceeds the theoretical limit of ∼20 s^-2^ at this field strength. We also plotted our experimentally determined relaxation times on simulated T_1_ and T_2_ curves over a range of τ_c_ and O^2^ (Figure 5f,g). At 5 °C, many helix and helix-adjacent residues possess O^2^ > 1.0 regardless of the simulated τ_c_ (Figure 5f). Taken together, this indicates the presence of exchange relaxation (R_ex_) occurring on the microsecond-second timescale. In the presence of R_ex_, the measured effective R_2_ rate (R_2_^eff^) is equal to R_2_ + R_ex_ which explains the elevated R_1_×R_2_ products at 5 °C [39]. We then simulated the effect of R_ex_ values on the R_1_xR_2_ product to estimate the magnitude of chemical exchange (Figure S6a). If we assume a highly-structured N_H_ O^2^ = 0.86, a commonly estimate for structured regions [37], then R_ex_ would be in the range of 12-42 s^-1^ at 5 °C when for a τ_c_ = 9-14 ns (Figure S6a). We conclude these additional exchange contributions reflect peptide:micelle binding kinetics although additional experiments would be needed to determine the on/off exchange rate.

Conversely, at 35 °C the R_1_×R_2_ values do not surpass the ∼20 s^-2^ limit and thus don’t appear to include significant R_ex_ contributions (Figure 5d and S5). It’s likely the increased temperature pushed the on/off exchange kinetics faster into the microsecond timescale, reducing R_ex_ contributions[37]. Further, the helical and helical adjacent residues N376-R392 cluster to τ_c_ ∼ 8 ns in a region with order parameter less than 1.0, however order parameters for several resonances are quite close to 1.0 (Figure 5g). Again, if we assume a highly structured N_H_ O^2^ = 0.86 and a τ_c_ of 8 ns, then the R_ex_ contributions at 35 °C are between 0.5 and 4 s^-1^ with a median position at ∼2.3 s^-1^ (Figure S6b). The tight clustering of nearly all helix residues to a highly ordered position at 8 ns on the T_1_/T_2_ plot (figure 5g) also supports minimal anisotropy. Taken together, intermediate timescale motions are reduced compared to at 5 °C but they are not eliminated completely.

### Determination of the H8 tilt and azimuthal angles relative to the DH_7_PC micelle

Published NTS1 structures containing an intact H8 indicate the amphipathic helix is oriented approximately parallel to the membrane surface [19-21, 26, 40-42]. We next sought to determine the orientation of the [*U*-^15^N,^13^C]-pNTS1(H8-Ctail) peptide relative to the micelle in solution at near physiologic temperatures using NMR. Respondek and colleagues showed that the paramagnetic relaxation enhancement (PRE) of ^1^H T_1_ values in alpha helices follows a wavelike pattern from which both tilt (τ; the angle between the helix axis and the micelle surface) and azimuthal (ρ; rotation of the helix) angles can be derived [43]. pNTS1(H8-Ctail) ^1^H_N_ T_1_ relaxation was measured in an inversion-recovery type sequence by appending a 180° ^1^H hard pulse and variable relaxation delay period to the beginning of a ^13^C-HSQC (Figure 6a). Plotting these ^1^H_N_ T_1_ relaxation times on a per residue basis produced a wavelike pattern that repeated every three to four residues (Figure 6b, red line).

**Figure 6.**
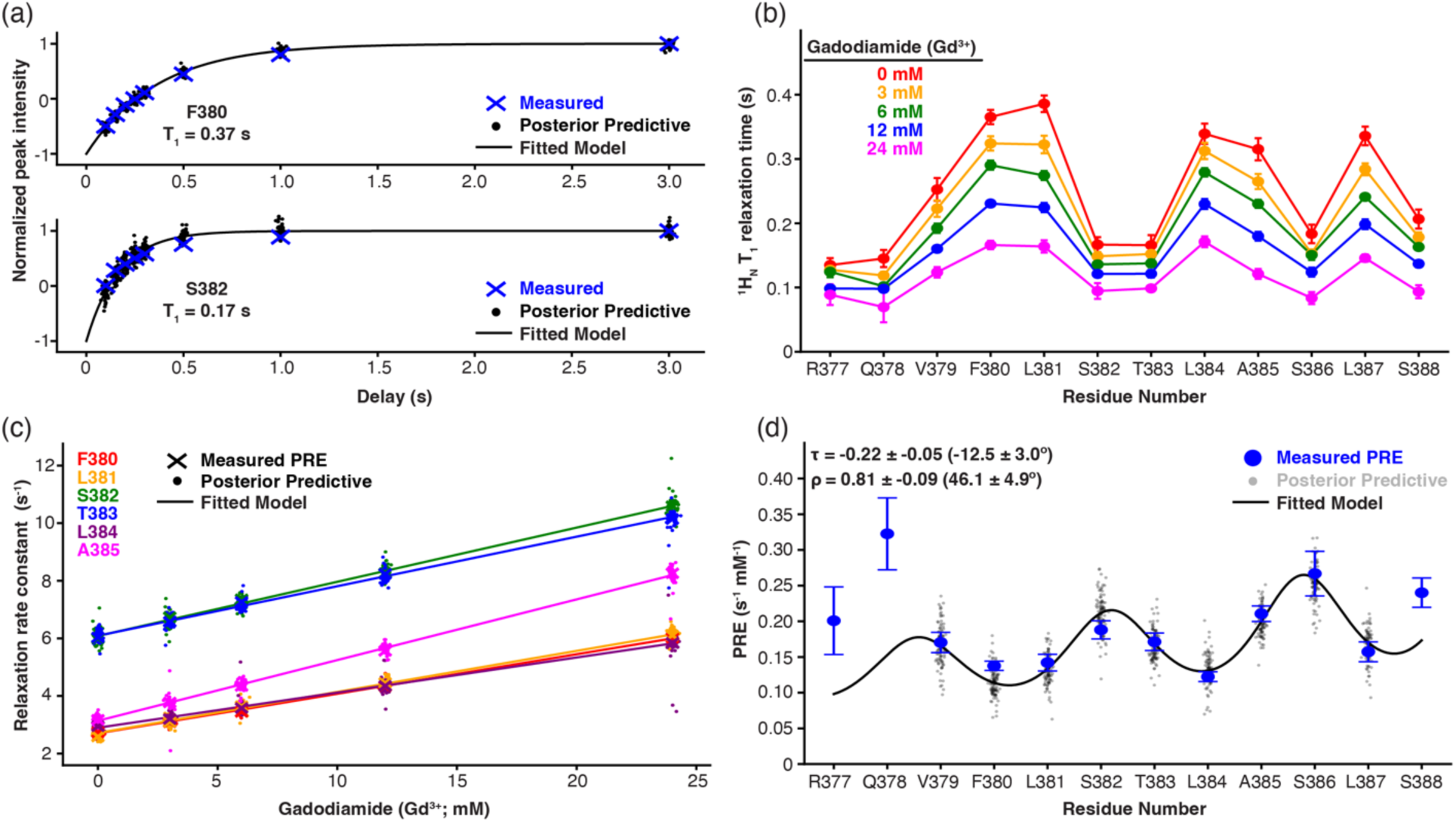
PRE of ^1^H_N_ T_1_ relaxation yields the pNTS1(H8-Ctail) helix orientation relative to the DH_7_PC micelles. a.) The peak intensity of F380 and S382 as a function of the inversion recovery delay serve as representative fits of ^1^H_N_ T_1_ relaxation times measured on pNTS1(H8-Ctail) in 12 mM (d26)-DH_7_PC at 35 °C. b.) Residue-specific ^1^H_N_ T_1_ relaxation times plotted for increasing gadodiamide concentrations. c.) The relaxation rate constants for several representative residues plotted as a function of gadodiamide concentration. The slope of each linear fit is the PRE. d.) PRE values of ^1^H_N_ nuclei plotted as a function of residue number. The paramagnetic relaxation wave equation for a micelle-embedded helix was fitted from V379-L387 yielding the indicated tilt (τ) and azimuthal (ρ) angles in radians (degrees).

We then used a water-soluble gadolinium(III)-based chelate named Gd-diethylenetriaminepentaacetic acid bis(methylamide), also known as Gd(DTPA-BMA) *or* gadodiamide, as a paramagnetic relaxation enhancing agent. As gadodiamide cannot permeate membrane mimetics, it produces a strong distance-dependent relaxation rate enhancement on ^1^H nuclei.[44] Titration of gadodiamide into a [*U*-^15^N,^13^C]-pNTS1(H8-Ctail) sample containing 12 mM (d26)-DH_7_PC produced dose-dependent decreases in ^1^H_N_ T_1_ relaxation (Figure 6b). Plotting these ^1^H_N_ R_1_ relaxation rates as a function of gadodiamide concentration yields a straight line with a slope that determines the PRE (Figure 6c). No chemical shift perturbations were detected up to 24 mM gadodiamide confirming the absence of non-specific interactions (data not shown). When these PRE values are plotted for each residue, they produce a wavelike pattern between residues V379 and L387 that reflects the amphipathic nature of H8 immersed in the (d26)-DH_7_PC micelle (Figure 6d, blue dots). The residue-dependent PRE values were then fitted to a wavelike function that assumes a micelle-immersed helix with the geometry of 3.6 residues per turn, a pitch of 1.5 Å per residue, and a 1.95 Å radius for H_N_ atoms [43]. The estimated tilt angle of -12.5 ± 3.0° corresponds to a helix orientation that’s near parallel to the micelle surface with the negative sign indicating the C-terminus is closer to solvent than the N-terminus. The azimuthal angle is measured assuming the points closest to and furthest from the micelle are 0° and 180°, respectively. The estimated azimuthal angle of 46.1 ± 4.9° means that F380, L384, and L387 are pointed approximately to the center of the micelle. Elevated PRE values near the H8 boundaries may indicate increased dynamics where even a minor population with higher solvent exposure could artificially skew values because their 1/d^3^ distance dependance.

## DISCUSSION

The helix 8 and C-tail of GPCRs are comparatively lesser studied receptor regions despite their importance in cell surface trafficking, activation, and transducer interactions.[5-16] Our study explores the structural and dynamic properties of the isolated rat NTS1 helix 8 and C-tail in solution. We demonstrated that a hydrophobic membrane-like environment is required to stabilize helicity within the helix 8 region of the GPCR, while the C-tail remains generally unstructured. Interestingly, not all micelles are equivalent – LMNG is considerably less efficacious than related DM and DDM detergents despite its superior stabilization of the transmembrane region. The Watts group previously analyzed the rat NTS1 H8 utilizing EPR spectroscopy, molecular dynamics simulations, and circular dichroism.[4] Their CD analysis of the isolated helix 8 (no C-tail) also concluded that helicity requires a detergent or lipid mimetic. Our combined NMR analysis identified a stable helix from F376-R392 with a break at P389-G390 followed by a flexible stretch. Similarly, the Watts group recorded EPR spectra at 170K (-103.15°C) and 295K (21.85°C) on NTS1 with sequentially spin-labeled residues 374 to 393 reconstituted in brain polar lipid (BPL) extract liposomes; they found two helical stretches separated by residues F380-L381 in the apo receptor that became contiguous upon agonist binding [4]. The peptide construct used in our study does not contain any transmembrane (TM) domain yet forms a single helical structure in DH_7_PC. It’s been suggested for other receptors [10, 12, 45-47] that ligand binding could affect H8 dynamics although all agonist and inverse-agonist bound NTS1 structures contain a single, unbroken helix. Our T1, T2 and PRE NMR data suggest regions adjacent to the H8 boundaries may form more transient helical structure. Similarly, no regular structure was discernable following P389 in Watts and colleagues’ EPR data [4].

Figure 7 summarizes the proteomicelle complex size and orientation at scale (Figure 7a,b). Both CD and NMR experiments support a 1:1 peptide:micelle molar stoichiometry and a helical length of 2.6 nm assuming it is 17 residues from 376-391. Helix 8 lies generally parallel to the micelle surface with a tilt angle of -12.5° (Figure 7a) and an azimuth angle of 46° that orients the hydrophobic residues (F376^8.50^, F380^8.54^, L381^8.55^, L384^8.58^, L387^8.61^) towards the micelle and hydrophilic residues (Q378^8.52^, S382^8.56^, S386^C8.60^) to the solvent. (Figure 7b, c). The negative tilt angle indicates the N-terminus is oriented deeper in the micelle surface plane than the C-terminus. This is the opposite orientation from both NTS1 structures and molecular dynamics (MD) simulations; the simplest explanation is that our study focuses on an isolated peptide lacking the TM domain. Although this negative tilt angle may reflect the broader range of H8 orientations accessible in a full-length receptor consistent with previous MD observations.

**Figure 7.**
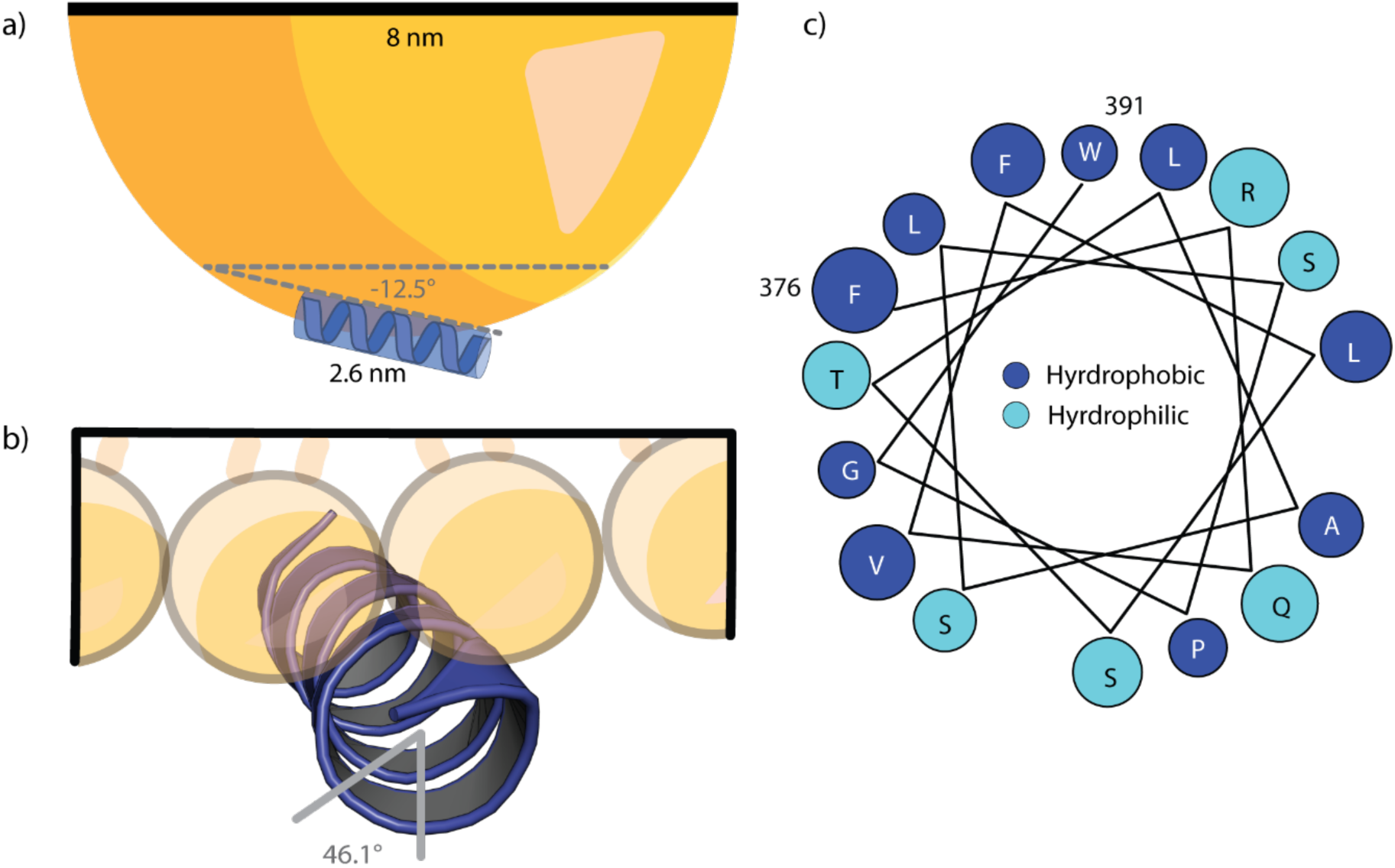
NTS1 helix 8 orientation within DH_7_PC micelle. a.) Model of NTS1 helix 8 tilt angle relative to the DH7PC micelle calculated from PRE data. b.) Model of NTS1 helix 8 azimuth (rotational) angle relative to the DH7PC micelle calculated from PRE data. c.) Helical wheel projection of NTS1 helix 8 based on the predicted helix 8 from calculate PRE values.

While the PRE calculations allowed for estimation of the helix orientations they do not permit a quantitative calculation of the insertion depth into the micelle. The Watts group used spin labeled POPC lipids to estimate the insertion depth of helix 8 into the membrane. The greatest increase in EPR order parameters when helix 8 was added was seen in carbon position 5 in the POPC outer leaflet indicating helix 8 primarily interacts with the lipid headgroups. This is in line with the predicted amphipathic nature of the helix and qualitatively with the observed PRE effects on our helix 8. Additionally, the MD models for isolated helix 8’s at varying lengths (374-385 and 373-393) show that the N-terminal half of the helix appears more stable than the C-terminal half. Interestingly, simulations of the full receptor showed a fully stable helix in BPL-like membrane, but only a partially stabilized helix in POPC alone. Similar to our CD data in various detergents where LMNG is less efficient at stabilizing helix 8, this suggests differences in the membrane mimetic can alter helix 8 architecture and stability. Interestingly, out of the rat NTS1 structures with helix 8 in the sequence, constructs in POPC/POPG nanodiscs, monoolein lipid with cholesterol, and nonylglucoside/octylglucoside/decylglucoside/dodecylglucoside (NG/OG/DG/DDG) detergents with cholesterol produced a structured helix 8.[19-21] Conversely, rat NTS1 structures in LMNG with cholesterol and glyco-diosgenin did not produce a structured helix 8.[48] However, these structures contain a scFv16 antibody which may be involved in the lack of helix 8 structure. All human NTS1 structures have a structured helix 8 and most of these are in LMNG supplemented with cholesterol, glyco-diosgenin, diC8-PtdIns(4,5)P2, and/or OG.

Finally, C386 and C388, known palmitoylation sites, were mutated to serines to avoid dissulfide formation. MD simulations on the dopamine D2 receptor indicated that H8 palmitoylation drives the helix further into the lipid membrane [49]. It’s not unreasonable to suspect these reversible modifications can influence, or be influenced by, receptor activation or transducer binding [50]. Simultaneous palmitoylation of the two NTS1 C-terminal cysteines was shown to be necessary for mitogenic signaling [27]; although individual mutations were not tested. Our PRE data indicates C388 is oriented closer to the membrane surface than C386 leading to questions surrounding the degree of palmitoylation and variable effects on receptor structure/function/dynamics. Further research is needed to establish how palmitoylation may stabilize H8 and regulate signaling.

## METHODS

### Reagents

Thermo Scientific™ HisPur™ Ni-NTA Resin used in the IMAC purification step was purchased from Fisher Scientific (#88221). Prepacked SP Ion Exchange column used in the pNTS1(H8-Ctail) purification was purchased from GE (#17-5247-01). Zorbax 300SB-C8 Semi-Preparative Reverse Phase HPLC Column was purchased from Agilent (#880995-206).

### pNTS1(H8-Ctail) Peptide Plasmid Construct

The DNA fragment encoding NTS1 Helix 8 and the C-tail (373-423, C386S, C388S), HRV 3C protease site (ETLFQGP), muGFP, and 10x-Histidine tag from previously characterized functional variant enNTS1[51] was cloned into a pET28a vector containing a *Saccharomyces cerevisiae* Smt3 with an N-terminal 6x-Histidine tag. The resulting expression vector contained an open reading frame encoding an N-terminal 6x-Histidine tag, followed by Smt3, the NTS1 Helix 8 + C-tail, HRV 3C protease site, a GGGSGGGS linker, muGFP, and a 10x-Histidine tag. The final sequence for the purified construct used in NMR experiments was: SANFRQVFLSTLASLSPGWRHRRKKRPTFSRKPNSMSSNHAFSTSATRETLFQ

### 15N,13C pNTS1(H8-Ctail) Expression

The pNTS1(H8-Ctail) plasmid was transformed into BL21(DE3) *Escherichia coli* cells and plated on LB agar supplemented with 50 µg/mL kanamycin and 1% (w/v) glucose at 37 °C overnight. Liquid LB media starter cultures were supplemented with 50 µg/mL kanamycin and 1% (w/v) glucose then inoculated with several colonies and incubated overnight at 37 °C and 220 RPM. The cells were used to inoculate 2L of H2O-based M9 media supplemented with 3 g L^-1^ of [U-^13^C_6_] d-glucose (Cambridge Isotope Laboratories, CIL) and 2 g L^-1^ of [^15^N] ammonium chloride (Cambridge Isotope Laboratories, CIL) as the sole carbon and nitrogen source, respectively. Cultures were incubated at 37 °C and 220 RPM until reaching an OD600 ≅ 0.5 at which point the temperature was dropped to 25 °C. Once each culture reached an OD600 ≅ 1, they were induced with 1 mM IPTG and incubated for ∼ 6 h at 25 °C and 220 RPM. The cultures were harvested via centrifugation at 5,000 g and cell pellets stored at -80 °C.

### 15N, 13C pNTS1(H8-Ctail) Purification

To prevent proteolytic degradation and avoid aggregation, the initial purification steps were done using 6M guanidine hydrochloride buffers. Cell pellets were solubilized in a *GdHCl solubilization buffer* (100 mM HEPES, 400 mM NaCl, 6M GdHCl, 20 mM imidazole, pH 8.0) supplemented with 0.2 mM PMSF. The solution was incubated on ice for 30 minutes before being subjected to sonication on ice: 5 minutes processing time (10 s on, 20 s off) at 35% maximum amplitude. A syringe fitted with an 18-gauge needle, bent at the tip, was used to further lyse and homogenize the solution. The solution was then centrifuged at 24,424 RCF for 45 minutes. The supernatant containing the pNTS1(H8-Ctail) construct was then collected and incubated with Ni-NTA resin equilibrated with *GdHCl solubilization buffer* at 4 °C for 45 minutes. Following resin incubation, the solution was placed into a gravity column to allow flow through of unbound protein. The Ni-NTA resin was then washed with *Wash #1* (25 mM HEPES, 400 mM NaCl, 6M GdHCl, 20 mM Imidazole, pH 8.0) followed by *Wash #2* (25 mM HEPES, 400 mM NaCl, 20 mM Imidazole, pH 8.0). This second wash step also serves to remove GdHCl and refold the Smt3 and muGFP fusion proteins on the column. Following GdHCl removal in *Wash #2*, the construct was eluted with *elution buffer* (25 mM HEPES, 200 mM NaCl, 500 mM Imidazole, pH 8.0) and incubated with 3 mg of ULP1 protease and 3 mg of HRV 3C precision protease for 6 hours at 4 °C to cleave Smt3 and muGFP respectively. The cleavage reaction was then diluted 10-fold in *SP equilibration buffer* (20 mM HEPES pH 8). The dilution was loaded onto an equilibrated 5 mL SP cation-exchange (CEX) column via a GE AKTA Pure system run at 4 mL/min flow rate. The SP CEX column was washed with *SP wash buffer* (20 mM HEPES, 400 mM NaCl, pH 8) until the AU280 stabilized. An initial elution was collected with *SP elution #1* (20 mM HEPES, 2M NaCl, pH 8). Likely due to the high isoelectric point of the pNTS1(H8-Ctail) peptide (∼12.4), a large portion of the peptide remained bound to the SP column despite the initial high salt elution. *SP elution #2* (100 mM HEPES, 400 mM NaCl, 6M GdHCl, pH 8.0) was found to completely remove the remaining peptide from the column. The two elutions were pooled and injected in several runs onto an Agilent C8 Semi-Preparative Reverse Phase HPLC Column. A 40-minute gradient of 32% to 42% acetonitrile with 0.2% TFA was applied to the column. MALDI-TOF mass spectrometry was used to identify and confirm purity of pNTS1(H8-Ctail) fractions. Fractions containing the pure peptide were pooled and subjected to lyophilization and then redissolved in water. The dissolved peptide was added to a 1 mL Float-A-Lyzer G2 Dialysis Device from Repligen with a 0.5-1 kDa cutoff. The peptide was initially dialyzed against 500 mL of water for >2 hours at room temperature with gentle stirring. This process was repeated with 500 mL of *dialysis buffer* (20 mM NaPi, 150 mM NaCl, pH 6.8) D_2_O and DSS were added resulting in the peptide being in a final *NMR buffer* (20 mM NaPi, 150 mM NaCl, 10% D_2_O, 200 µM DSS, pH 6.8).

### Circular dichroism

For CD measurements, the synthetic pNTS1(C-tail) peptide or synthetic pNTS1(H8-Ctail) peptide were diluted to a final concentration of 0.2 mg/mL in a *CD buffer* (10 mM NaPi pH 6.8). Peptide concentration was measured using absorption at 280 nm. CD spectra were recorded between 190 and 260 nm on a Jasco J-715 Spectropolarimeter using a 1-mm quartz cuvette, a scan rate of 50 nm/min, and a data integration time of 1 s. The collected spectra were corrected for buffer contribution accounting for the appropriate detergent and temperature before being converted to molar ellipticity. Spectra for the synthetic pNTS1(C-tail) peptide were collected at 5, 15, 20, 30, 45, 60, 70, 80, 90 °C. Additionally, spectra were collected at 20 °C with 3 mM DH_7_PC and 12 mM DH_7_PC. Spectra for the pNTS1(H8-Ctail) peptide were collected at 5, 15, 20, 25, 30, 35, 45, 60, 70, 80, 90 °C. For the pNTS1(H8-Ctail) spectra were also collected with 0.5, 1.5, 3, 6 mM, and 12 mM DH_7_PC at 20 °C. Additional detergent comparisons were collected at 20 °C with 0.05%, 0.1%, 0.3%, and 0.5% DM; 0.05%, 0.1%, 0.3%, and 0.5% DDM; 0.005%, 0.05%, 0.1%, and 0.5% LMNG.

### NMR spectra assignments experiments

NMR spectra were collected on a Bruker AVANCE NEO II 14.1 T (Indiana University – Bloomington) spectrometer equipped with a HCN cryogenic probe. DSS was present in all samples to reference spectra. ^1^H-^15^N HSQC spectra were recorded for 450 µM [^15^N, ^13^C] pNTS1(H8-Ctail) in *NMR buffer* at 5, 10, 15, 20, 25, 30, and 35 °C. HNCA, HNCOCA, HNCO, HNCACO, HNCACB, and HNCOCB experiments were used to assign backbone amides in the ^1^H-^15^N HSQC at 5 °C. These assignments were transferred to other temperatures by tracking chemical shift perturbations. Similarly, ^1^H-^13^C HSQC spectra were recorded for 450 µM [^15^N, ^13^C] pNTS1(H8-Ctail) in *NMR buffer* at 5, 10, 15, 20, 25, 30, and 35 °C. Additional 15N-TOCSY, 15N-NOESY, (H)CC(CO)NH, and H(CCCO)NH spectra were collected to assign carbon and proton resonances in the ^1^H-^13^C HSQC. Again, assignments were transferred to other temperatures by tracking chemical shift perturbations.

### (d26)-DH_7_PC NMR titration and assignment

Tail deuterated (d26)-DH_7_PC powder was dissolved in *NMR buffer* at a stock concentration of 500 mM. ^1^H-^15^N HSQC and ^1^H-^13^C HSQC spectra were recorded for 450 µM [^15^N, ^13^C] pNTS1(H8-Ctail) in *NMR buffer* at 5°C with addition of 0, 0.5, 1.5, 3, 6, 10, and 12 mM (d26)-DH_7_PC. At 12 mM (d26)-DH_7_PC, a temperature series of ^1^H-^15^N HSQC and ^1^H-^13^C HSQC spectra was collected at 5, 10, 15, 20, 25, 30, and 35 °C. ^1^H-^15^N HSQC and ^1^H-^13^C HSQC spectra assignments at 12 mM and 35 °C generated using chemical shift tracking from previous conditions and collection of HNCA, HNCO, (H)CC(CO)NH, and H(CCCO)NH.

### Chemical shift perturbation mapping

Chemical shift perturbations for [^15^N, ^13^C] pNTS1(H8-Ctail) were calculated using ^1^H-^15^N HSQC spectra comparing conditions 5°C and 35°C at 0 mM (d26)-DH_7_PC; 5°C and 35°C at 12 mM (d26)-DH_7_PC; 0 mM and 12 mM (d26)-DH_7_PC at 5°C; 0 mM and 12 mM (d26)-DH_7_PC at 35°C. Euclidean distance calculations were measured based on PPM position with N chemical shifts being weighted by a factor of 0.142 to H chemical shifts.

### Peak height comparison

Absolute peak heights for [^15^N, ^13^C] pNTS1(H8-Ctail) residues in ^1^H-^15^N HSQC spectra with 12 mM (d26)-DH_7_PC at 5 °C and 35°C were measured and the peak height differences between the two temperatures were calculated. These values were then plotted for each residue.

### TALOS-N secondary structure prediction

TALOS-N was used to predict phi and psi backbone torsion angles and estimate secondary structure of [^15^N, ^13^C] pNTS1(H8-Ctail) using the measured N, NH, Cα, Cβ, CO, Hα, and Hβ chemical shift values. These values from the peptide at 5°C/0 mM (d26)-DH_7_PC and 35°C/12 mM (d26)-DH_7_PC were submitted to the TALOS-N web server (https://spin.niddk.nih.gov/bax-apps/software/TALOS-N/) and results were analyzed using jRAMA+.

### ^15^N Relaxation and ^1^H Paramagnetic relaxation enhancement (PRE) with gadodiamide paramagnetic probe

^15^N relaxation data (T_1_ and T_2_) were collected at 600 MHz on a Bruker AVANCE NEO II (Indiana University – Bloomington). Started pseudo3D interleaved pulse sequences were used with delay times of 0.1, 0.2, 0.3, 0.4, 0.5, 0.6 s for T_1_ and 0.0157, 0.0314, 0.0470, 0.0627, 0.0786, 0.0941 s for T_2_. Relaxation data were analyzed in SPARKY and relaxation parameters extracted using the Sparky ‘rh’ command.

To measure ^1^H PRE, gadodiamide powder was dissolved in *NMR buffer* at a stock concentration of 500 mM. T_1_ recovery time spectra with delay times of 0.10, 0.15, 0.20, 0.25, 0.30, 0.50, 1.0 and 3.0 seconds were recorded for 350 µM [^15^N, ^13^C] pNTS1(H8-Ctail) in *NMR buffer* plus 12 mM (d26)-DH_7_PC with addition of 0, 3, 6, 12, and 24 mM gadodiamide at 35 °C. Spectra were recorded as a series of ^15^N HSQCs with an appended 180° ^1^H pulse at the start of the sequence, followed by the T_1_ recovery delays above. T_1_ relaxation times were estimated for peptide residues by measuring change in peak height relative to increasing delay times for each gadodiamide concentration. First, all peak height values were normalized to the final delay time (3 seconds) peak height. Normalized data was then fitted to the following model using Bayesian Parameter Estimation:

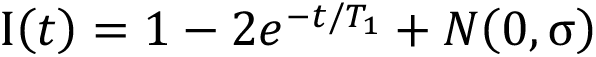

where 𝐼 is the peak heights, 𝑡 is the delay time. In this model the values of 𝐼(𝑡) ranges from -1 at 𝑡 = 0 and 1 at 𝑡 = ∞, representing full inversion and full recovery. After 3 seconds of delay recovery (our longest delay) we appeared to have achieved very close to full recovery (Figure 6a). Extracted parameters are 𝑇_1_, the inversion recovery (T_1_) estimate and σ, the standard deviation of a normal distribution centered at zero, 𝑁(0, σ). In this model, σ is a measure of the error of our data to the fitted model. For ^1^H spins measured in the putative helix the magnitude of σ did not exceed 0.1 or 5% of the range from -1 to 1, indicating our data is well described by the fitted model. This can be seen by the posterior predictive data points in Figure 6a (black dots), which are predictions of data from the fitted model at the time delays recorded.

T_1_ values at the gadodiamide concentrations listed above were then fitted to a straight-line model to extract the PRE as the gradient of the line. The following Bayesian Parameter Estimation model was used:

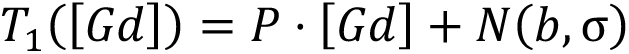

Where 𝑇_1_([𝐺𝑑]) is the measured T_1_ value above at concentration of Gadodiamide [𝐺𝑑]. Extracted parameters are 𝑃, the PRE, 𝑏, the offset T_1_ value at a Gd concentration of zero and 𝜎 is a measure of the error of our data to the fitted model. Values of σ were always lower than 5% of the highest 𝑇_1_([𝐺𝑑]) value, indicating the above linear model describes our data well out to 24 mM gadodiamide. This can be seen with the posterior predictive data points generated from our fitted model in Figure 6c.

The PRE values were used to estimate the orientation of the helix with respect to the micelle by using the following model derived by Respondek *et al* [43] for a helix buried away from a PRE agent, inside a micelle, modified with a noise factor:

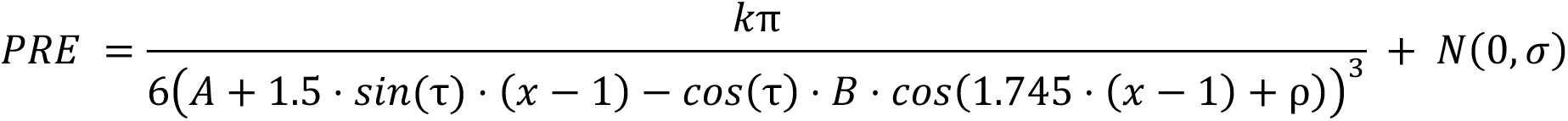

where 𝑥 is the number of the amino acid in the sequence of the helix, starting at the N-terminus, 𝐴 is the immersion depth of the helical axis at the first residue, 𝐵 is the radius of the helix for the atoms undergoing PRE (1.95 Å for 1H_N_). The parameters estimated from data are τ, the tilt angle, ρ, the azimuth or the rotation angle of the helix, 𝑘, a constant to account for a combination of proportionality constants in the system to do with the strength of the PRE effect from the gadodiamide and finally, 𝜎, to account for noise in the data during Bayesian Parameter Estimation. The estimation of σ was 0.019 which is ∼7% of the highest PRE value recorded in the helix at ∼0.27 s^-1^mM^-1^ for S386, showing our fitted model is in good agreement with the data (see Figure 6d).

## Supplementary Information

**Figure S1.**
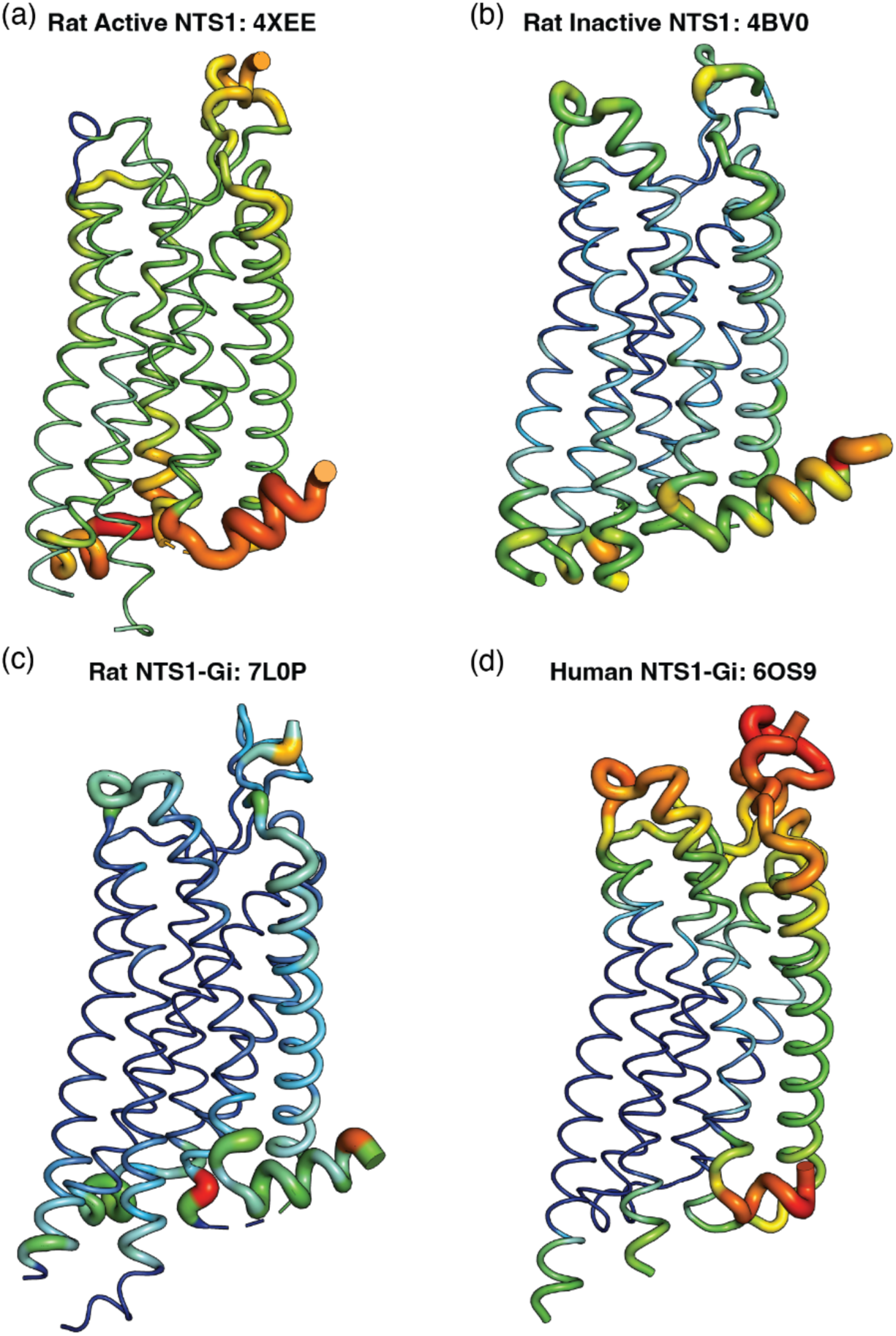
The H8 region typically possesses the highest crystallographic B-factors across NTS1 structures. PyMOL B-factor putty representations for several NTS1 structures: (a.) active-state rat NTS1 (PDB 4XEE), (b.) inactive-state rat NTS1 (PDB 4BV0), (c.) heterotrimeric canonical Gi-bound rat NTS1 (PDB 7L0P), and (d.) heterotrimeric canonical Gi-bound human NTS1 (PDB 6OS9).

**Figure S2.**
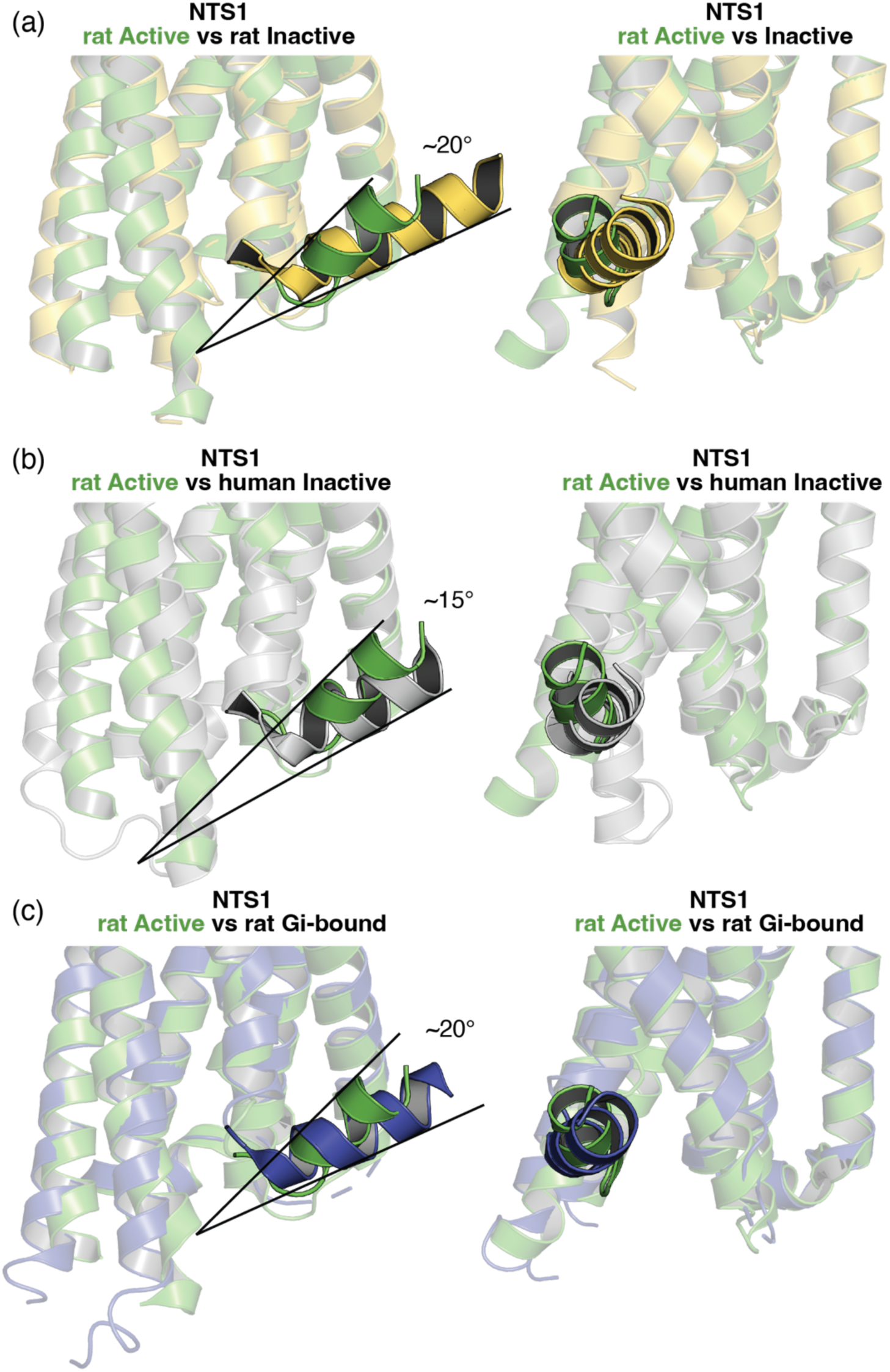
Comparison of helix 8 orientation across NTS1 structures. Alignment of active-state rat NTS1 (green; PDB: 4XES) with (a.) inactive-state rat NTS1 (yellow; PDB: 4BWB), (b.) inactive-state human NTS1 (gray; PDB: 7UL2), and (c.) heterotrimeric Gi-bound rat NTS1 (blue; PDB: 7L0P). Approximate difference in the helix 8 tilt angle between structures is shown.

**Figure S3.**
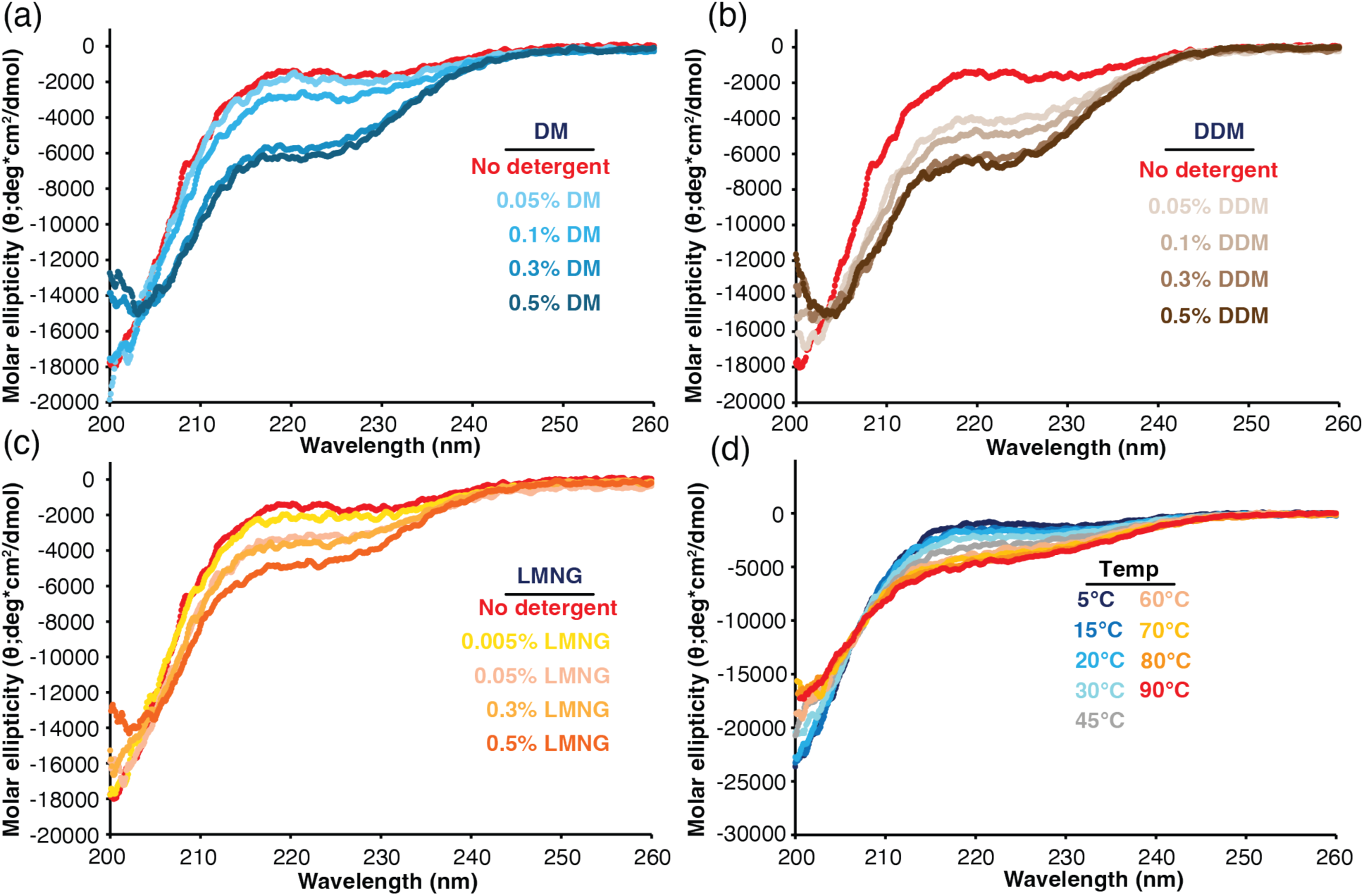
CD spectra of NTS1 peptides as a function of temperature or detergent concentration. Overlay of pNTS1(H8-Ctail) CD spectra at 20 °C as a function of increasing (a.) DM, (b.) DDM, or (c.) LMNG detergent concentration. d.) Overlay of pNTS1(Ctail) peptide CD spectra as a function of temperature.

**Figure S4.**
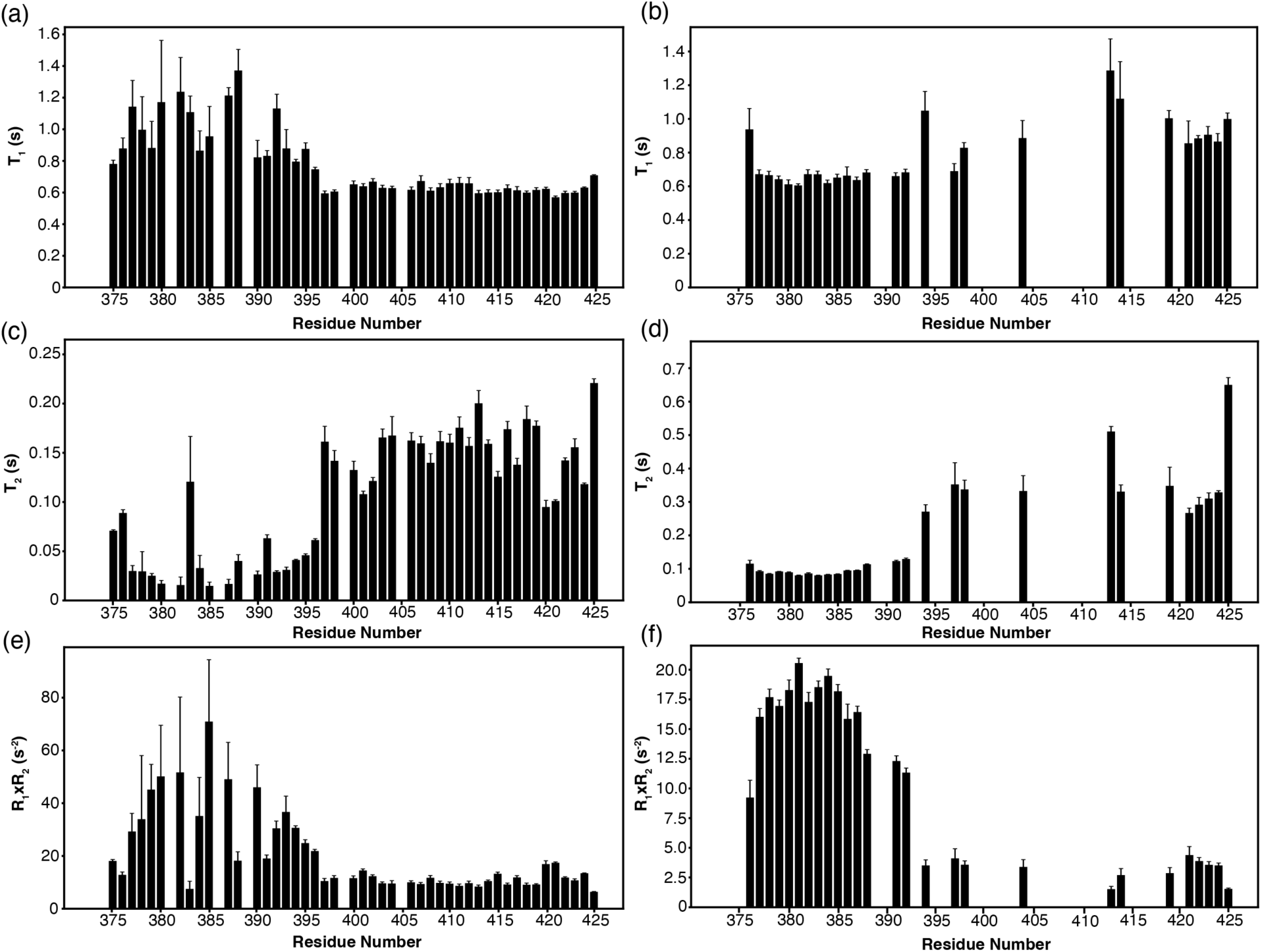
pNTS1(H8-Ctail) ^15^N T_1_ and T_2_ relaxation at 5 °C and 35 °C. pNTS1(H8-Ctail) ^15^N T_1_ values were calculated by fitting peak heights to a two parameter exponential decay function at (a.) 5 °C and (b.) 35 °C. pNTS1(H8-Ctail) ^15^N T_2_ values were calculated by fitting peak heights to a two parameter exponential decay function at (c.) 5 °C and (d.) 35 °C. pNTS1(H8-Ctail) ^15^N R_1_xR_2_ values with propagated errors at (e.) 5 °C and (f.) 35 °C.

**Figure S5.**
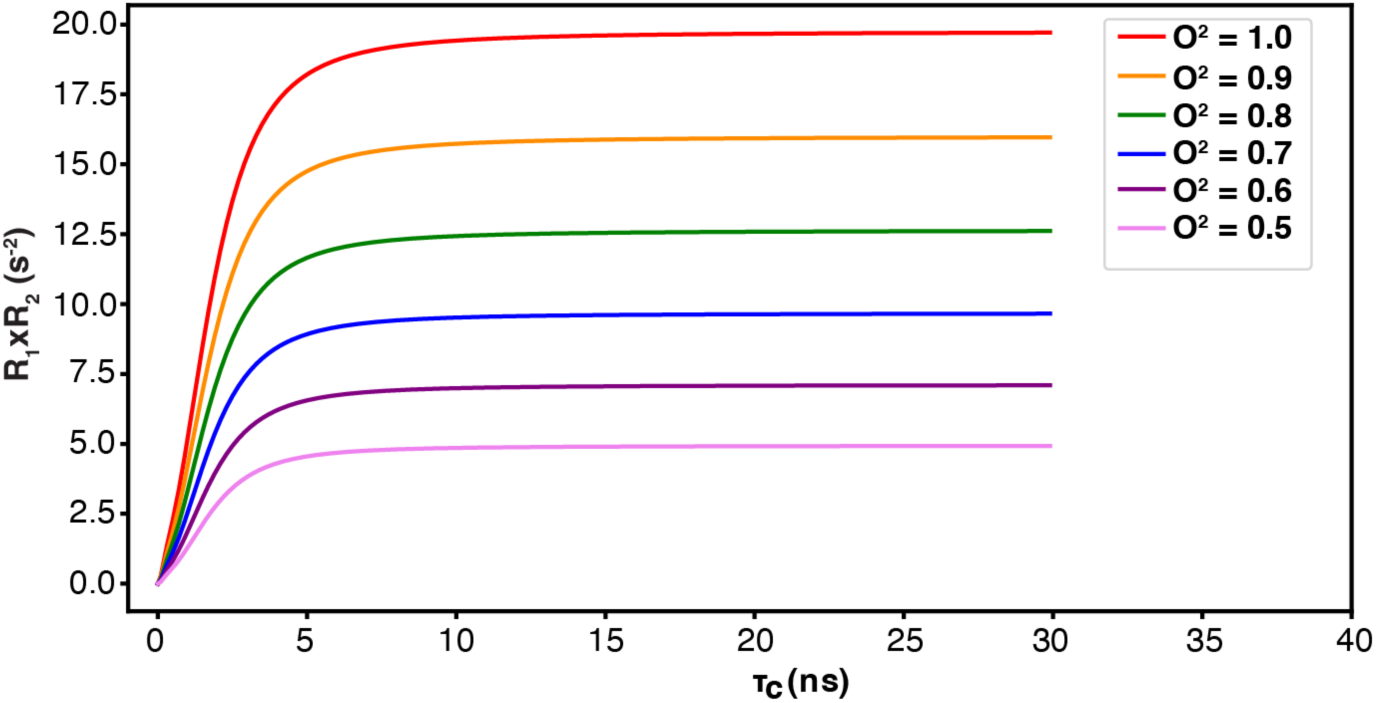
Simulated R_1_×R_2_ values at 600 MHz magnetic field strength. Generalized order paramaters (O^2^) ranging from 0.5 to 1.0 were used in simulations of R_1_ and R_2_. R_1_xR2 values were calculated from the simulated R_1_ and R_2_ values with τ_𝑐_ values in the range 0 to 30 ns and plotted for each O^2^ value.

**Figure S6.**
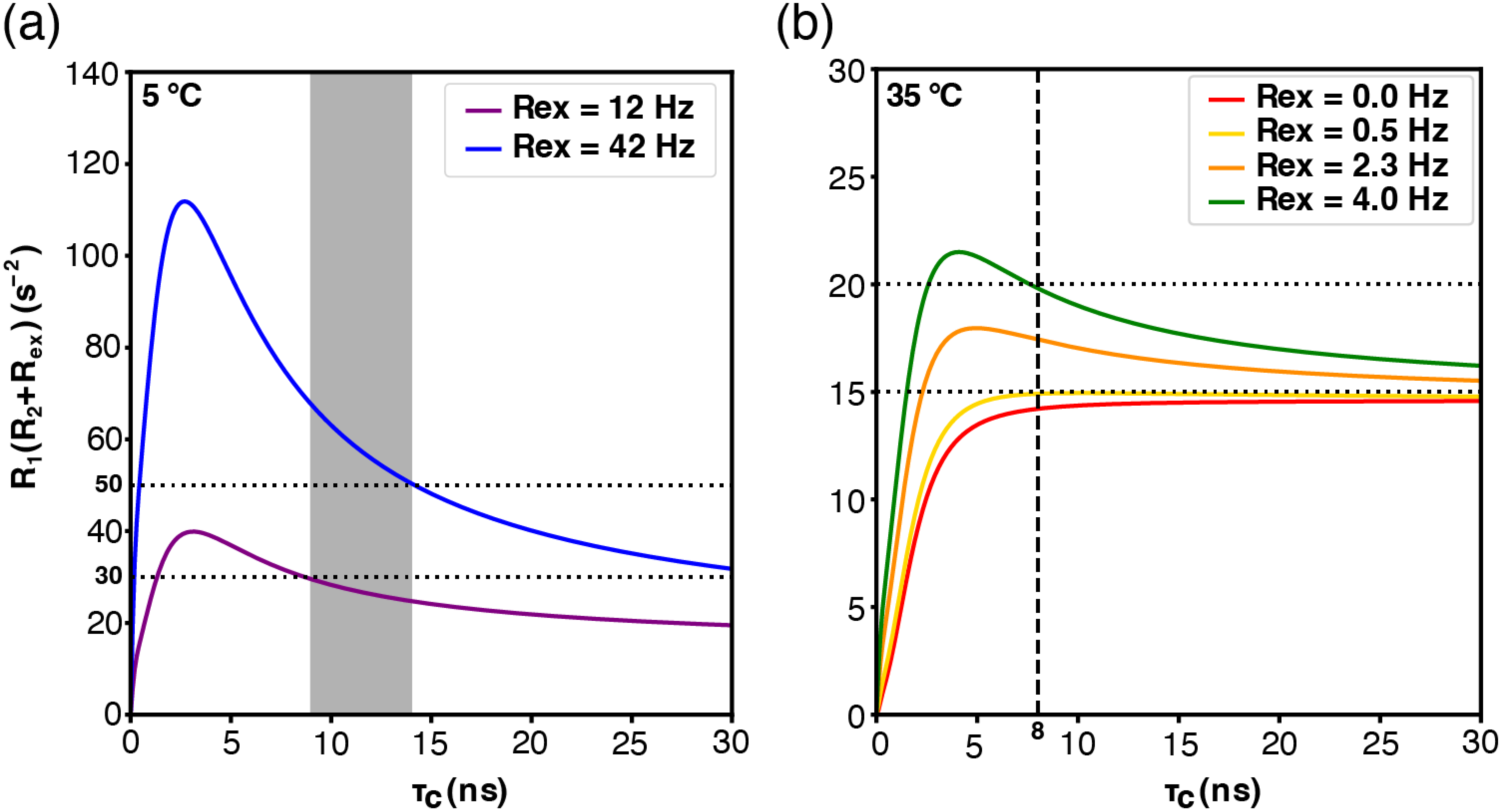
Simulated R_1_xR_2_ values versus 𝛕_𝒄_ including contributions of R_ex_ to R_2_ for rigid structure (O^2^ = 0.86). (a.) The range of measured R_1_xR_2_ values at 5 °C (30-50 s^-2^) are indicated with horizontal dashed lines. The grey vertical bar reflects the estimated τ_c_ ranges for the micelle. Additional R_ex_ contributions of 12 and 42 Hz are plotted in purple and blue, respectively. (b.) The range of measured R_1_xR_2_ values at 35 °C (15-20 s^-2^) are indicated with horizontal dashed lines. The dashed vertical line indicates the 8 ns estimated τ_c_ value. R_ex_ contributions of 0.0, 0.5, 2.3 and 4.0 Hz are plotted in red, yellow, orange and green, respectively.

